# MPXV Infects Human PBMCs in a Type I Interferon-Sensitive Manner

**DOI:** 10.1101/2024.09.16.613292

**Authors:** Laure Bosquillon de Jarcy, Dylan Postmus, Jenny Jansen, Julia Melchert, Donata Hoffmann, Victor M. Corman, Christine Goffinet

**Affiliations:** Institute of Virology, Campus Charité Mitte, Charité - Universitätsmedizin Berlin, Charitéplatz 1, 10117 Berlin, Germany; Berlin Institute of Health, Berlin (BIH), Anna-Louisa-Karsch-Str. 2, 10178 Berlin, Germany; Department of Infectious Diseases and Respiratory Medicine, Charité - Universitätsmedizin Berlin, Charitéplatz 1, 10117 Berlin, Germany; Friedrich-Loeffler-Institut, Südufer 10, 17493 Insel Riems, Germany; German Center for Infection Research, associated partner Charité, Berlin, Germany; Liverpool School of Tropical Medicine, Liverpool, L35QA UK

**Keywords:** MPOX, MPXV, monkeypox, viral dissemination, PBMCs, tropism

## Abstract

MPOX virus (MPXV), formerly known as monkeypox virus, led to a rapidly evolving pandemic starting May 2022, with over 90,000 cases reported beyond the African continent. This pandemic outbreak was driven by the MPXV variant Clade IIb. In addition, Clade I viruses circulating in the Democratic Republic of Congo (DRC) are drawing increased attention as cases constantly rise and Clade Ib, first identified in 2023, is now co-circulating with Clade Ia and seems to exhibit enhanced human-to-human transmissibility. While most infected individuals display a self-limiting disease with singular pox-like lesions, some endure systemic viral spread leading to whole-body rash with risks for necrosis, organ loss, and death. Intra-host dissemination and cellular tropism of MPXV are largely unexplored in humans. To establish a potential susceptibility of circulating immune cells to MPXV, we exposed human PBMCs from healthy donors *ex vivo* to a currently circulating MPXV clade IIb virus isolate in absence and presence of IFN-α2a. qPCR of DNA extracted from cell lysates, but less from supernatants, revealed increasing MPXV DNA quantities that peaked at five to six days post-exposure, suggesting susceptibility of PBMCs to infection. IFN-α2a pretreatment markedly reduced the quantity of MPXV DNA, suggesting that infection is sensitive to type I IFNs. Plaque assays from supernatants showed that infection gave rise to *de novo* production of infectious MPXV. In virus-inclusive scRNA-sequencing, monocytes, cycling NK cells and regulatory CD4^+^ T-cells scored positive for viral RNA, suggesting that these are the MPXV-susceptible cell types within the human PBMC population. Analysis of differentially expressed genes displayed a pronounced downregulation of expression pathways driving innate immunity in MPXV-infected cells, a well-established feature of poxviral infection. Pretreatment of PBMCs with current antivirals Cidofovir and Tecovirimat resulted in reduced amounts of viral antigen production and of released infectivity, suggesting suitability of the human PBMC infection model as a platform for evaluation of current and future antivirals and justifying trials to investigate Cidofovir and Tecovirimat as drugs reducing intra-patient viral spread. Together, our data suggest that human PBMCs are productively infected by MPXV which is accompanied by significant modulation of the cellular milieu. Our results have the potential to illuminate aspects of intra-host propagation of MPXV that may involve a lymphohematogenous route for replication and/or intra-host dissemination.

## Background

MPOX is an emerging zoonosis causing fever and painful rash with pox-like lesions in infected humans. The causative virus belongs to the family of the *Orthopoxviridae* and consists of two clades: Clade I, endemic in the Congo Basin, being the more virulent clade with mortality rates in humans reported up to 10%, while clade II, endemic in West Africa, leads to comparably milder disease courses of MPXV infection (Bunge et al.). Transmission occurs via ingestion of body fluids from infected individuals, inhalation of infectious aerosols or close skin contact with infectious pox-like lesions. Sexual transmission of the virus is speculated, as infectious MPXV particles could be detected in semen of infected individuals and pronounced genital lesions were observed (Lum et al. 2022). MPOX has also been discussed to be transmitted in a sex-associated manner, not absolutely requiring sexual intercourse for transmission but rather depending on close skin to skin contact. In May 2022, MPXV clade II first spread to a rapidly evolving pandemic with over 90,000 cases worldwide, representing the largest outbreak of MPOX beyond the African continent ever recorded (Liu et al. 2023). The pandemic-causing circulating clade II strain was subsequently designated as clade IIb (Americo, Earl, and Moss 2023). While the number of cases in the MPOX Clade IIb outbreak has plummeted in most parts of the world due to rising awareness of risk groups and vaccination, the virus continues to circulate, with currently most cases being reported in South East Asia and the Western Pacific Region (“Multi-Country Outbreak of Mpox, External Situation report#35-12 August 2024”). Furthermore, the recent surge in Clade I MPOX cases in the Democratic Republic of Congo (DRC) has attracted considerable attention. The DRC is currently experiencing its largest recorded outbreak of Clade I MPOX. As of September 2024, the CDC has reported over 33,799 confirmed and suspected cases, along with more than 1,000 deaths with fatality rates from 1.4% – >10%, mostly affecting young children (McQuiston et al. 2024). MPOX clade I has evolved in two different strains: Clade Ib circulates in the eastern DRC and neighbouring countries, where the outbreak predominantly affects adults and spreads mainly through sexual contact. Clade Ia circulates in areas of the DRC where MPOX is endemic, with the disease mostly affecting children and spreading through multiple modes of transmission (Vakaniaki et al. 2024). In response, the WHO has declared a public health emergency of international concern on 14th of August 2024 (“WHO Director-General Declares Mpox Outbreak a Public Health Emergency of International Concern”).

Infection by the pandemic-causing circulating MPXV clade IIb provokes a mostly self-limiting course of disease. However, in some individuals, the virus spreads systemically and leads to a whole-body rash with massive inflammation, causing necrosis, organ loss, and even death (Patel et al. 2023). These severe courses of disease have mostly, but not solely, been reported in immunocompromised patients, where the virus dissemination seems to be more excessive while other infected individuals display locally delimited viral dissemination (Miller et al. 2022). Since intra-host MPXV dissemination and cellular tropism have been studied insufficiently in humans, the underlying cause for these varying clinical observations remains poorly understood. Available data are primarily derived from other poxviruses, such as vaccinia virus, where monocytes appear to be an important target cell type (Lum et al. 2022), and from experimental infection of non-human primates (NHP), which suggest that monocytes are a susceptible cell type within the PBMC population (Johnson et al. 2011). For MPXV, tropism in human PBMCs remains unexplored, resulting in crucial knowledge missing to develop targeted therapeutic strategies preventing extended intra-host dissemination and severe courses of disease. From the point of view of pandemic preparedness, understanding poxvirus pathogenesis will be essential to deal with future outbreaks of MPXV and further Orthopoxviruses.

Adding to the complexity, efforts to develop antiviral compounds against poxviruses have almost come to a standstill since smallpox was eradicated in 1979. As a result, specific therapeutic options for the treatment of MPOX have been minimally studied in humans (Lum et al. 2022). Three compounds, not all of them licensed for MPOX treatment, are currently available: Tecovirimat, an inhibitor of VP37 membrane protein on the surface of orthopoxviruses, impairs budding of *de novo*-produced virions. Survival after MPXV infection was improved by Tecovirimat treatment in NHPs (Huggins et al. 2009). In humans, the data on the benefits of Tecovirimat against MPOX is very limited (Siegrist and Sassine 2023). One clinical study showed no significant difference in the clinical course of MPXV Clade IIb infections between patients who received Tecovirimat and those who received symptomatic therapy only (Ouyang et al. 2024). Another study conducted in the DRC also failed to demonstrate any clinical benefit from Tecovirimat in infections with MPOX clade I (Lenharo 2024). Cidofovir, a nucleotide analogon impairing MPXV DNA polymerase, has shown efficacy in humans infected with molluscum contagiosum (Siegrist and Sassine 2023). Brincidofovir, a Cidofovir analogon, displays improved cellular uptake due to its enhanced lipophilicity and showed efficacy in prairie dogs infected with MPXV (Siegrist and Sassine 2023). However, clinical effectiveness of Cidofovir and Brincidofovir in humans with MPOX is solely documented in singular case reports and has not yet been analysed in clinical studies. As these three antiviral compounds are also very limited in availability, treatment for MPOX is usually restricted to symptomatic and supportive therapy for the vast majority of cases. Therefore, the identification, development and preclinical evaluation of antiviral compounds against MPXV is urgently needed.

## Methods

### Cells

VeroE6 cells (ATCC CRL-1586) were cultured in Dulbecco’s modified Eagle’s medium (DMEM) complemented with 10% heat-inactivated foetal calf serum (FCS), 1% penicillin– streptomycin (Thermo Fisher Scientific) and 2 mM L-glutamine (Thermo Fisher Scientific) at 37°C in a 5% CO_2_ atmosphere. Cell lines were monitored for the absence of mycoplasma.

Human PBMCs from anonymised healthy blood donors were isolated from 7 ml EDTA whole blood. Withdrawal of blood samples from healthy humans and cell isolation was conducted with approval of the local ethics committee (Ethical review committee of Charité Berlin, vote EA1/193/22 of 2022, Nov 15th). Samples were diluted 1:1 in PBS and centrifuged on Pancoll (Pan Biotech) for 30 min at 200 x *g*. PBMCs were washed with PBS and remaining erythrocytes were lysed with ACK-Lysis Buffer (8,29g NH_4_Cl, 1g KHCO_3_, 0,0367g EDTA, 600ml H_2_O), followed by PBS washing. After isolation, PBMCs were cultured in RPMI 1640 supplemented with 10% heat-inactivated FCS (Sigma Aldrich), 1% penicillin–streptomycin (Thermo Fisher Scientific), 2mM L-glutamine (Thermo Fisher Scientific), 1% non-essential amino acids (NEAA, Thermo Fisher Scientific) and 1% sodium pyruvate (NaP, Thermo Fisher Scientific). The experiments conformed to the principles of the WMA Declaration of Helsinki and the Department of Health and Human Services Belmont Report.

### Virus

MPXV clade IIb was isolated from a human specimen collected in the pandemic of 2022 in Berlin and propagated on Vero E6 cells and concentrated using Vivaspin® 20 columns (Sartorius Stedim Biotech). MPXV stocks were diluted in OptiPro serum-free medium complemented with 0.5% gelatine and PBS and stored at −80°C. Infectious titer was defined via plaque titration assay. Whole-genome sequencing was performed, and the resulting sequence has been deposited in the GISAID under accession number EPI_ISL_13890273 (Jones et al. 2022).

### MPXV Infection

Vero E6 cells were seeded in 24-well plates at densities of 3×10^5^ cells/ml in a total volume of 500 ul/well. Cells were infected with MPXV (MOI 0.00006, 0.0006 and 0.006) and spinoculated for 60 min at 800 x *g* followed by two hours of incubation at 37°C and 5% CO_2_. Afterwards, virus inoculum was removed and cells were washed two times with DMEM and supplied with fresh medium.

Human PBMCs were seeded in 96-well plates at densities of 1×10^5^ cells/ml in a total volume of 100ul/well. Cells were infected with MPXV (MOI 0.000006, 0,00006, 0,0006, 0,006) followed by spinoculation and incubation as indicated for Vero E6 cells. After incubation, cells were washed twice with RPMI and supplied with fresh medium. Where indicated, cells were pretreated with Interleukin-2 (IL-2, 20 nM, Merck) and Phytohemagglutinin (PHA, 2 ug/ml, Sigma Aldrich) for three days prior to infection and IL-2 treatment was continued until the end of the experiment. A subset of PBMCs was treated with IFN-α2a (500 IU/ml, Roche) overnight prior to infection and treatment was continued until the end of the experiment.

### Antiviral Compounds

A subset of PBMCs was pretreated with antiviral drugs 90 minutes before infection with MPXV. We used Tecovirimat (SIGA Technologies, New York, USA) and Cidofovir (Cayman Chemical, Michigan, USA) at a final concentrations of 5 μM (Grosenbach and Hruby 2019), (Frenois-Veyrat et al. 2022) and 17 μM (Andrei and Snoeck 2010), respectively. The compounds were readded after washing of the cells after infection and remained in the cell culture until the end of the experiment.

### Viral DNA Isolation and qPCR

For isolation of viral MPXV DNA, 300 μl of MagNA Pure 96 external lysis buffer (Roche, Penzberg, Germany) was added to 50 μl of supernatant or to dry cell pellets. All samples were heat-inactivated for 10 minutes at 70°C prior to export from the BSL-3. Isolation and purification of viral DNA was performed using the MagNA Pure 96 System (Roche, Penzberg, Germany) according to the manufacturers’ recommendations. Viral DNA was quantified using real-time PCR targeting duplicated G2R genes. Oligonucleotides 5′-GGAAAATGTAAAGACAACGAATACAG-3′ (forward primer), 5′-GCTATCACATAATCTGGAAGCGTA-3′ (reverse primer) and 5′-FAM-AAGCCGTAATCTATGTTGTCTATCGTGTCC-BHQ1-3′ (fluorescent probe) were used (Y. Li et al. 2010). Relative DNA levels were determined using the ΔΔCT method, with human *RNASEP* DNA (Applied Biosystems) as internal reference.

### Plaque Assays

Plaque assays were performed to determine the infectious titer in supernatants of MPXV-infected PBMCs at multiple time points. 2×10^5^ Vero E6 cells were seeded in a 24-well plate one day prior to infection. Cells were inoculated with 200 ul supernatant (1:200 to undiluted). After incubation for one hour at 37°C, supernatants were removed from the Vero E6 cells and 500 ul of overlay (1:1 mix of 2.4% avicel and 2×concentrated DMEM supplemented with 5% FCS, 2% NEAA, and 2% NaP) was added. After incubation for 72 hours, overlay was discarded and cells were fixed for 30 minutes in 6% PFA, then washed once with PBS and stained for 20 minutes with crystal violet solution. Infectious titer was calculated by division of the number of plaques by the respective inoculation volume and multiplied with the inverted dilution factor.

### Immunoblotting

Cell lysates were generated with 1xSDS sample loading buffer (Sigma-Aldrich, St. Louis, Missouri, USA). Proteins were separated on a 10% SDS-PAGE and transferred onto nitrocellulose using a semi-dry transfer system (Bio-Rad Laboratories, Hercules, California, USA). Membranes were blocked with 5% milk powder solution for one hour and incubated overnight with a polyclonal anti-orthopox rabbit serum (1:1000) (Czerny et al. 1994). Secondary antibodies conjugated to Alexa 680/800 fluorescent dyes were used for detection and quantification of expression by Odyssey Infrared Imaging System (LI-COR Biosciences Lincoln, NE, USA).

### Flow Cytometry

Cells were fixed in 4% PFA (Carl Roth) and permeabilised in 0.1% Triton X-100 (Thermo Fisher Scientific) in PBS before immunostaining with a polyclonal anti-orthopox rabbit serum (1:1000) (Czerny et al. 1994). Secondary antibodies conjugated to Alexa Fluor 488 or 647 (1:1000; Invitrogen) were used for detection. Flow cytometry analysis was performed using FACS Celesta with BD Diva Software (BD Biosciences) and FlowJo V10.8 Software (FlowJo).

### Single-Cell RNA-Sequencing

PBMCs from one healthy donor were isolated and infected with MPXV (MOI 0.0006) *ex vivo* or mock-infected. PBMCs were harvested three and five days post-infection for scRNA-seq.

Single-cell RNA-seq libraries were prepared with the 10× Genomics platform using the Chromium Next GEM Single Cell 3’ Reagent Kits v.3.1 following the manufacturer’s instructions. Quality control of the libraries was performed with the KAPA Library Quantification Kit and Agilent TapeStation. Libraries were sequenced on a HiSeq4000 using the following sequencing mode: read 1: 28 bp, read 2: 91–100 bp, Index i7: 8 bp. The libraries were sequenced to reach ∼20,000 reads per cell.

### Single-Cell RNA-Sequencing Data Analysis

FASTQ files from the sequencing protocol were processed using the Cell Ranger pipeline v 3.1.0 (10× Genomics) and further analysed using the Seurat v3.1.4 package (Butler et al. 2018) in R v3.6 (https://www.r-project.org/). Preprocessing of the data was performed using the recommended SCTransform procedure and the IntegrateData with PrepSCTIntegration workflows to eliminate batch effects. Gene expression values were normalised using the NormalizeData function and the “LogNormalize” setting. A comprehensive description of the code used in the analysis of data is available at https://github.com/GoffinetLab/MPXV_PBMC_study. Cell types were identified based on marker gene expression as outlined by the Seurat tutorial (https://satijalab.org/seurat/articles/pbmc3k_tutorial): B cells (CD3D^+^, MS4A1^+^), CD4^+^ T-cells (CD3D^+^, CD8A*), CD8^+^ T-cells (CD3D^+^, CD8A^+^), NK cells (CD3D^-^, CD8A^-^, NKG7^+^, GNLY^+^), monocytes (CD3D^-^, CD14^+^, FCGR3A^+^), dendritic cells (DCs, FCER1A^+^, CST3^+^), plasmacytoid dendritic cells (pDCs, LILRA4^+^). Additionally, activated CD4^+^ T-cells were identified by their expression of GZMA, GZMB or GZMK, while naïve CD4^+^ or CD8^+^ T-cells were defined as SELL^+^ (CD62L^+^) and memory subsets of both cell types as S100A4^+^. Regulatory T-cells were identified by elevated expression of IL2RA, IKZF2 and FOXP3, while NK cells undergoing mitosis (cycling) were defined by elevated expression of TOP2A. Reads aligning to the MPXV genome were identified by alignment to an MPXV Clade II (NC_063383.1, GenBank) reference using the same Cell Ranger pipeline. Mock-infected samples showed a negligible amount of reads aligned to the MPXV reference genome.

### Data Presentation and Statistical Analysis

If not stated otherwise, bar graphs indicate mean values and error bars indicate standard deviation. Graphs were generated using *Graph Pad Prism 9.1.2.* P-values < 0.05 were considered significant and labelled accordingly: P < 0.05 (*), P < 0.01 (**), or P < 0.001 (***); n.s. = not significant (≥ 0.05). Statistical overrepresentation analysis was performed with the list of DEGs harbouring p-values <0.05, gene set enrichment analysis (GSEA) was performed using the Pathway Panther Reactome database (Mi et al. 2019; Thomas et al. 2022). The results are described using the Fold Enrichment Score, indicating the degree of overrepresentation of a given gene set in the list of DEGs.

## Data Accessibility

The raw sequencing datasets generated in this study will be made available at the NCBI Gene Expression Omnibus upon publication and are currently available upon request.

## Results

### MPXV Infects PBMCs in a Type I Interferon-Sensitive Manner

To elucidate a possible blood immune cell tropism of MPXV, human PBMCs from three anonymous donors were exposed to MPXV at different MOIs (0.00006, 0.0006, 0.006, 0.06). Prior to infection, PBMCs were either mock-treated (FIG. 1A) or IL-2/PHA-stimulated (FIG. 1B), the latter resulting in a cell culture enriched in activated T-cells. In both types of PBMC cultures, MPXV DNA became detectable over time and peaked five to six days post-infection. DNA levels tended to be higher in IL-2/PHA-stimulated PBMCs, which may hint towards a proviral cellular milieu of activated T-cells. MPXV DNA quantities were reduced when cells were pretreated with IFN-α2a (500 IU/ml), suggesting sensitivity of viral DNA replication to type I IFN. Interestingly, IFN treatment-associated reduction of MPXV DNA levels was less effective at high MOIs, in line with the known saturability of IFN-induced antiviral factors by excess of viral antigens (FIG. 1). MPXV DNA was detectable in supernatants, albeit at lower amounts and showed a less pronounced increase over time if at all (Suppl. FIG. 1). This suggests a viral spread predominantly through cell-to-cell transmission rather than the release of infectious particles into the extracellular space, as known from other poxviruses such as variola virus, where infectious virus could be isolated infected blood cells, but not from cell-free plasma (Jahrling et al. 2004).

**FIG. 1.**
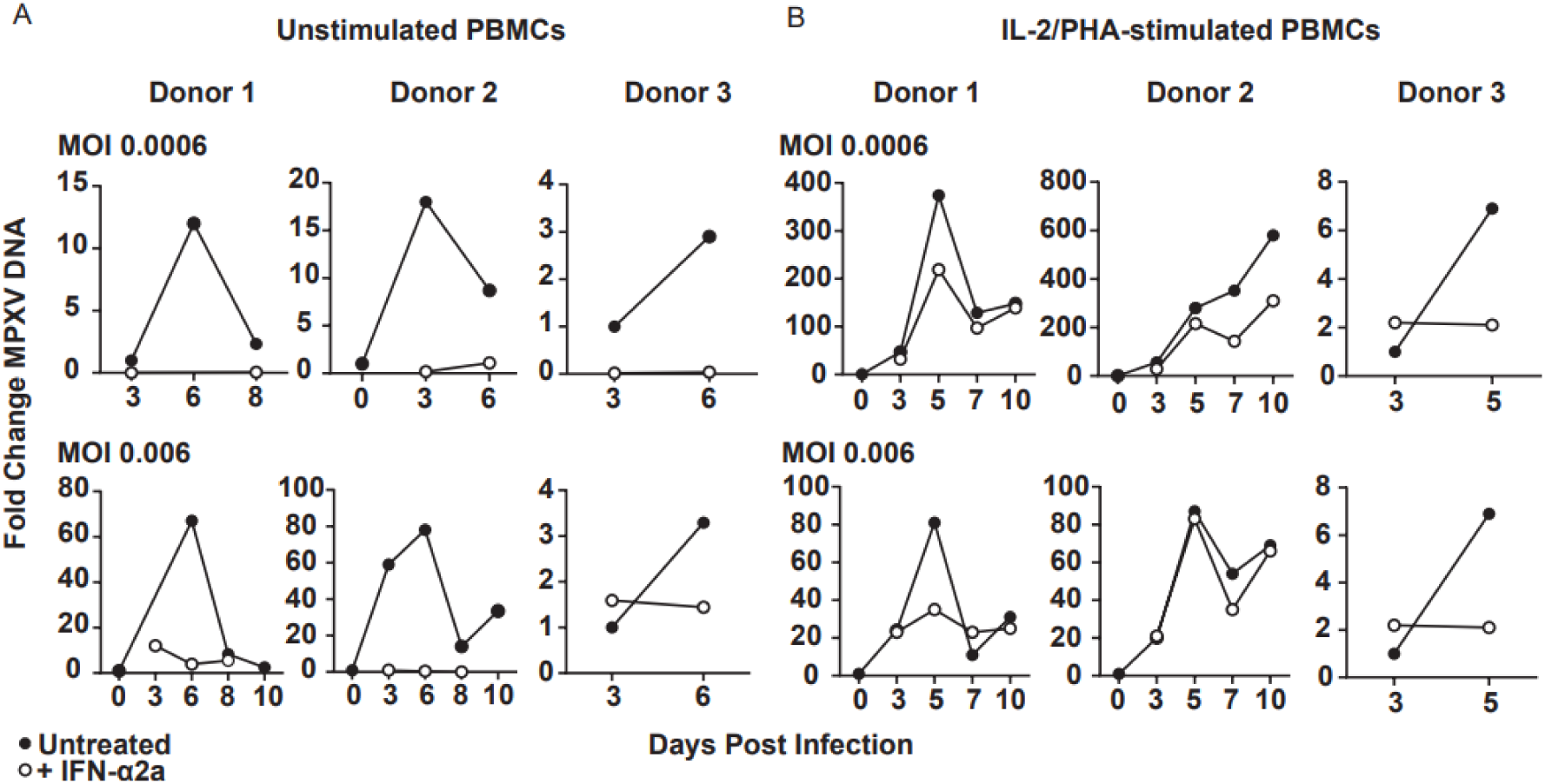
MPXV Infects PBMCs in a Type I Interferon-Sensitive Manner. A) PBMCs from three healthy donors were isolated, cultured and exposed to MPXV at increasing MOIs (determined on Vero E6 cells) in presence or absence of IFN-α2a (500 IU/ml) and harvested at multiple time points post-infection for quantification of cell-associated MPXV DNA by qPCR. Shown is the fold change of MPXV DNA relative to each donor’s MPXV DNA amount at the earliest time point and lowest MOI and was calculated via the ΔΔCT method using *RNASEP* as cellular gene. B) PBMCs from three healthy donors were isolated and cultured as described in A) but additionally stimulated with IL-2/PHA for three days prior to infection.

### MPXV Infection of Human PBMCs Results in Production of Infectious Virus Progeny

To determine whether MPXV infection of human PBMCs is productive or abortive, we transferred supernatants from *ex vivo* MPXV-infected PBMCs to Vero E6 cells for plaque assays which indicated the production of new infectious virions over time (FIG. 2). Titers were higher in IL-2/PHA-stimulated PBMCs than in unstimulated cells, suggesting that a population of expanding and activated T-cells is favourable to viral replication and/or release. In both stimulated and unstimulated PBMCs, viral titers gradually increased over time following infection. Notably, plaque formation was reduced when PBMCs were pretreated with IFN-α2a.

**FIG. 2.**
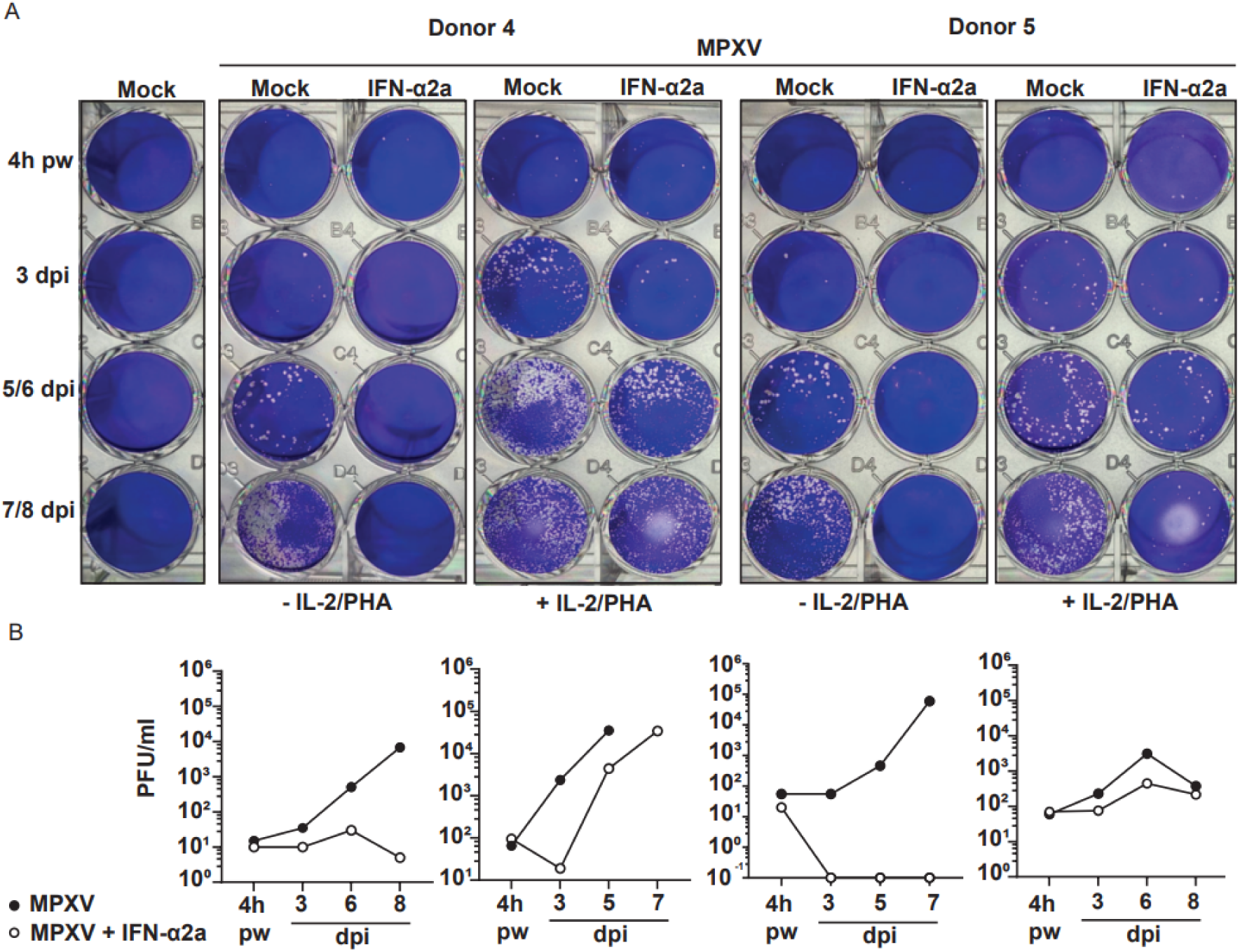
MPXV Infection of Human PBMCs Results in Production of Infectious Virus Progeny. A) Plaque assays were performed on Vero E6 cells using supernatants from MPXV-infected PBMC. PBMCs from two healthy donors were isolated, either mock- or pre-treated with IFNα (500 IU/ml) overnight, and infected *ex vivo* with MPXV (MOI 0.0006) and. Supernatants were harvested at indicated time points post infection, including one condition four hours post-infection (4h pw), which was harvested after removal of viral inoculum and washing of PBMCs with RPMI medium. Due to the limited sample volume, plaque counts were derived from a single well per dilution. B) Infectious viral titers in supernatants of MPXV-infected PBMCs shown in A.

### MPXV Infection of PBMCs Gives Rise to Viral RNA Positivity in Monocytes, Regulatory CD4^+^ T-Cells and NK Cells

To identify the target cell types of MPXV within the human PBMC population, we performed virus-inclusive single-cell RNA sequencing (scRNA-seq) on PBMCs from one healthy donor (Donor 4) infected ex vivo with MPXV. While mock-infected cells scored negative for viral RNA, as expected, monocytes, cycling NK cells and regulatory CD4^+^ T-cells displayed abundantly detectable MPXV RNA reads at day five post-infection (FIG. 3). Interestingly, at day three post-infection, only monocytes displayed notable quantities of viral RNA, suggesting that infection of this cell type occurs more rapidly and/or more efficiently compared to regulatory CD4^+^ T-cells and NK cells which scored positive only at day five post-infection. Examination of the viral transcriptional profile in these three cell types demonstrated extensive genome coverage, with expression of virtually all early, intermediate and late genes in monocytes and regulatory CD4^+^ T-cells, and many in cycling NK cells, at day five post-infection (Suppl. FIG. 2). This suggests that the MPXV replication cycle is entirely completed in these three cell types, consistent with our findings of a productive infection as evidenced by the plaque assays. Interestingly, individual viral genes were expressed in the other cell types, raising the question whether these cells might be infected incompletely or abortively, and/or infection requires more time to progress than the five days time window of our experiment.

**FIG. 3.**
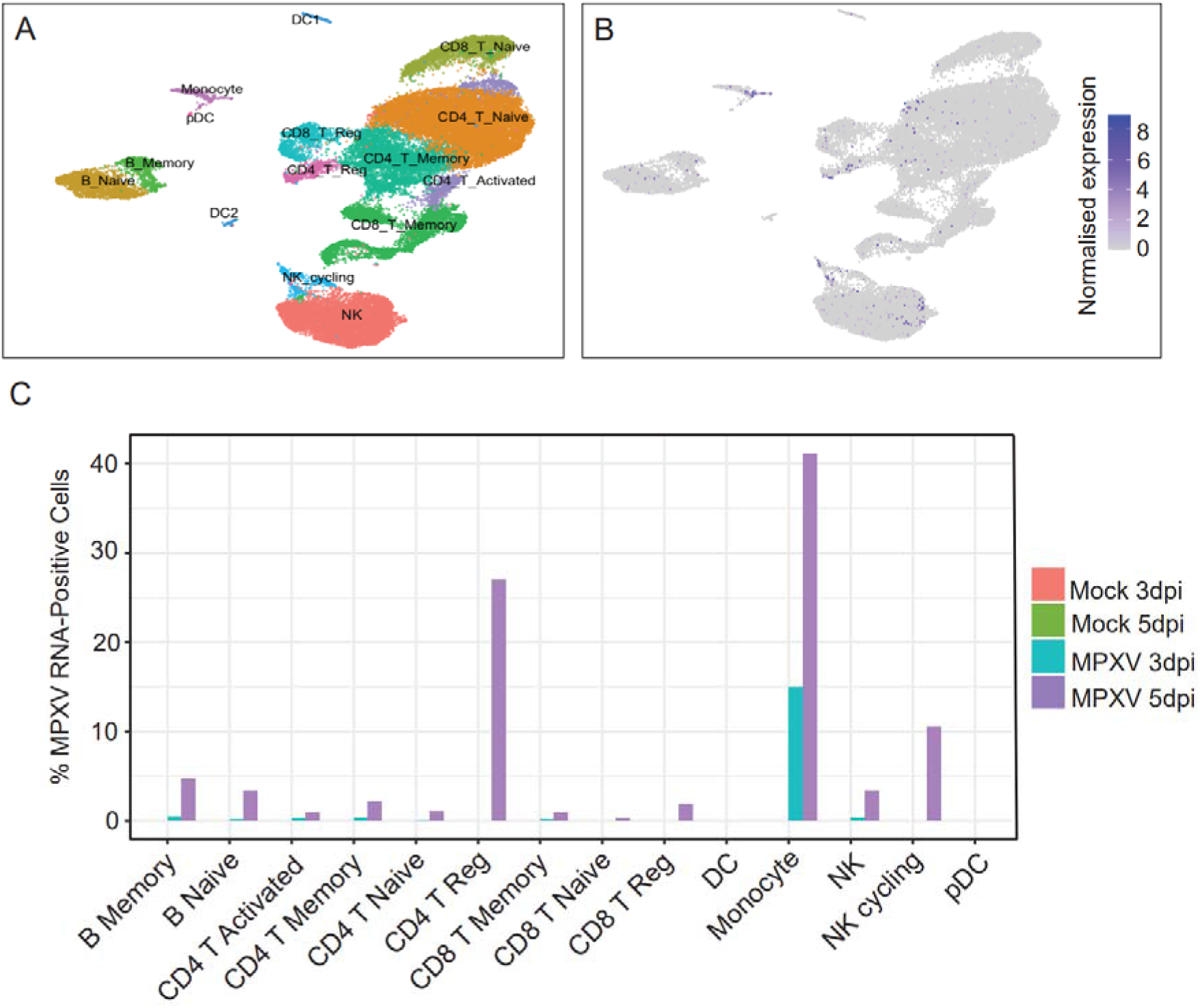
MPXV Infection of PBMCs Gives Rise to Viral RNA Positivity in Monocytes, Regulatory CD4^+^ T-Cells and NK Cells. PBMCs from one healthy donor (Donor 4) were infected with MPXV (MOI 0.0006) *ex vivo* or mock-infected. PBMCs were harvested three and five days post-infection for scRNA-seq. A) UMAP projection of PBMCs subjected to scRNA-seq three and five days following either mock or MPXV exposure coloured by cell type. B) UMAP plot showing log-normalised expression levels of MPXV RNA reads. C) Percentage of cells displaying ≥ ten viral RNA reads within each cell type or cell subset.

### Cell-Intrinsic Immune Responses are Downregulated in MPXV-Exposed and -Infected Monocytes, Regulatory CD4^+^ T-Cells and Cycling NK Cells

To investigate cell-intrinsic and cell type-specific responses to MPXV infection, we conducted differential gene expression (DGE) analysis. We compared gene expression between MPXV-exposed and mock-exposed PBMCs, as well as between MPXV RNA-positive and MPXV RNA-negative cells within the MPXV-exposed culture. Monocytes exhibited the highest number of DEGs (FIG. 4A), with many of these genes involved in innate immune responses and IFN-γ signalling (FIG. 4D). Notably, these genes were downregulated following MPXV exposure or infection (FIG. 4A-D). This downregulation included key components of the innate immune response such as *STAT1, JAK2, TRAC and GBP1, GBP2, GBP4, GBP5* (FIG. 4C). These findings are consistent with observations in other orthopoxviruses, which are known to suppress host antiviral innate immune responses (Hernaez and Alcamí 2024).

**FIG. 4.**
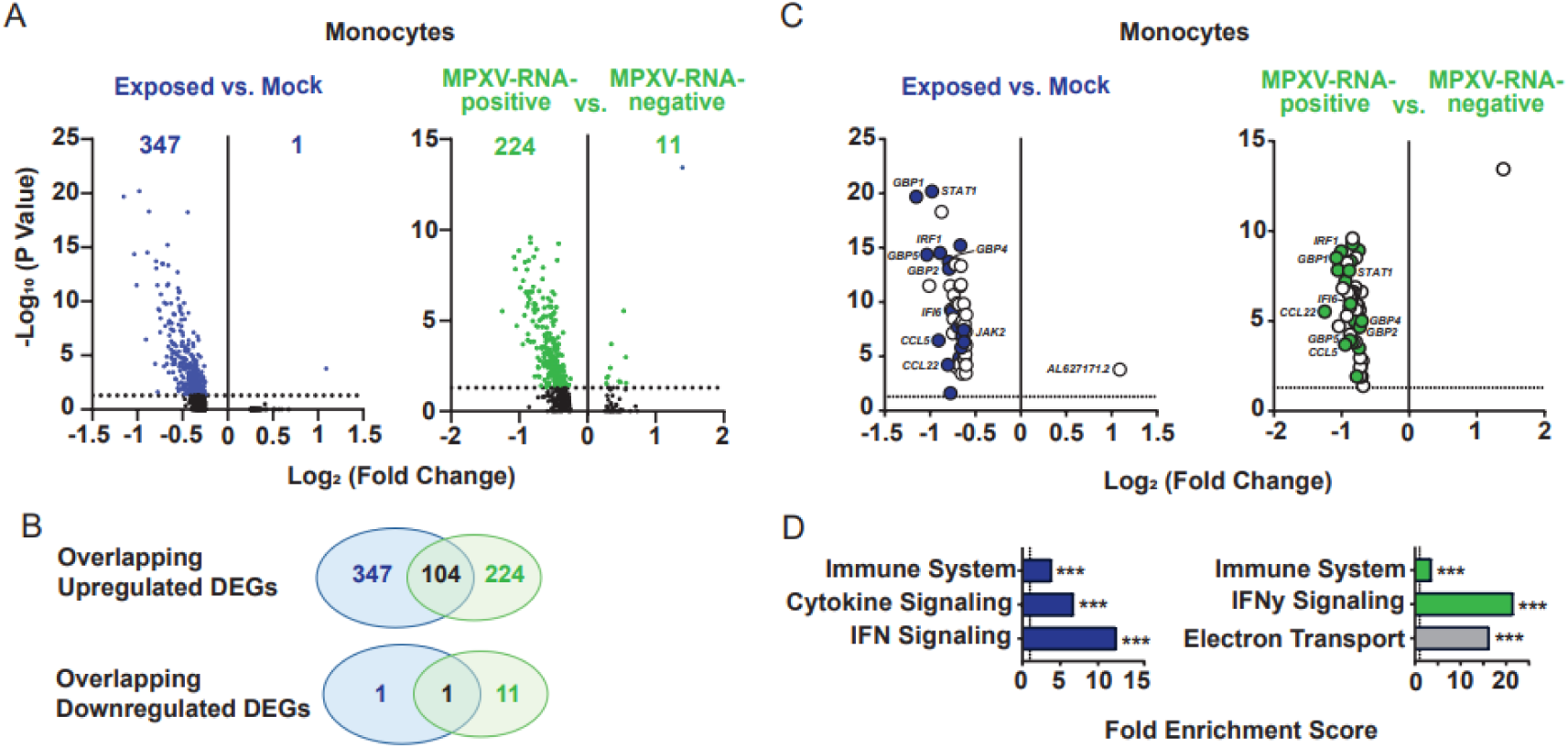
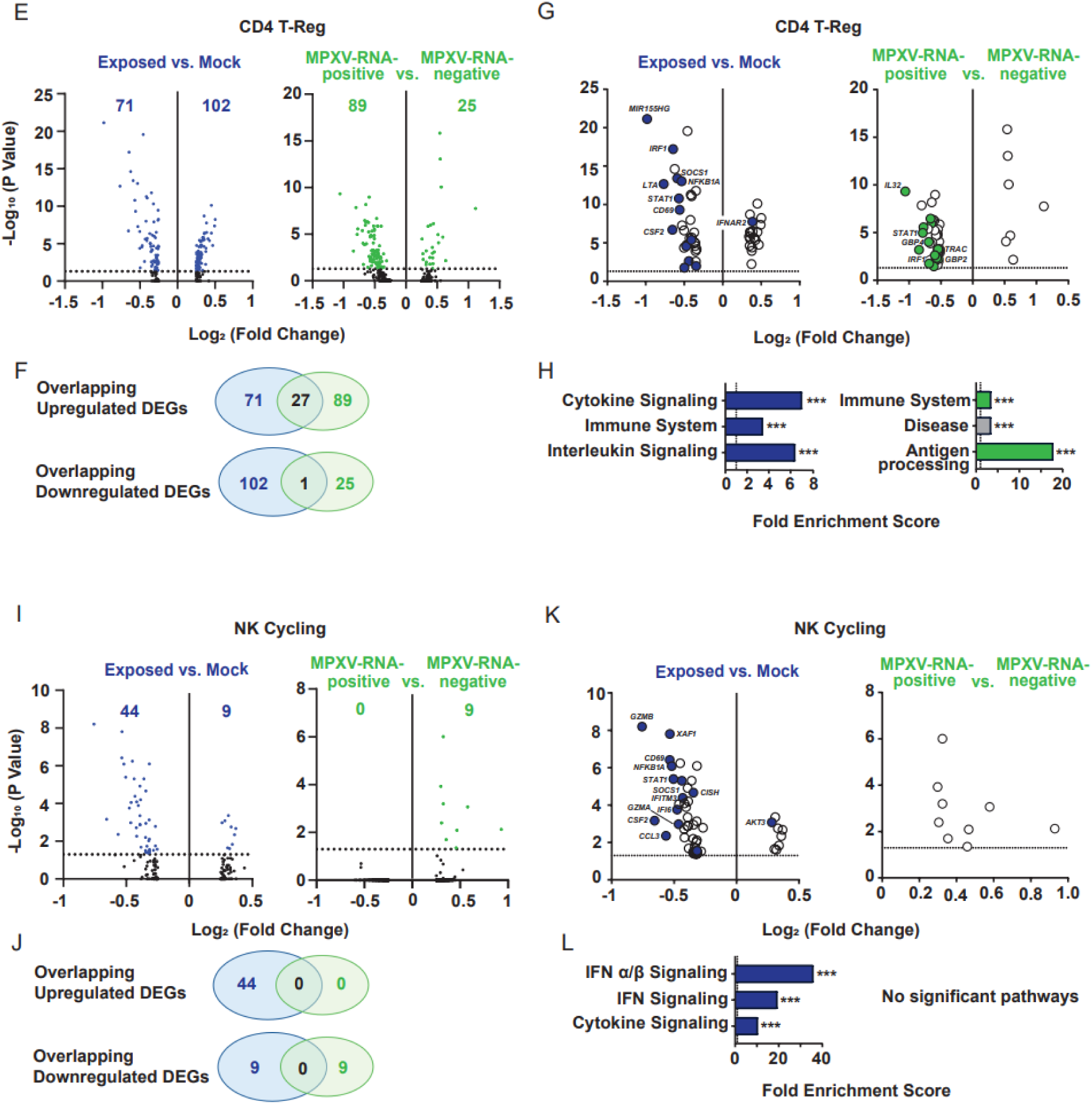
Cell-Intrinsic Immune Responses are Downregulated in MPXV-Exposed and - Infected Monocytes, Regulatory CD4^+^ T-Cells and Cycling NK Cells. The scRNA-seq data set shown in FIG. 3 was analysed for the expression profiles of human genes. A,E,I) Volcano plots showing Log_2_ Fold Change (< −0.25 and > 0.25) and statistical significance of DEGs for the contrasts MPXV-exposed vs. mock-exposed cells, and MPXV-RNA-positive cells vs. MPXV RNA-negative cells within the MPXV-exposed culture, for each target cell type or subtype at day five post-infection. Dotted line indicates a p-value of 0.05. Amount of significantly altered DEGs according to adjusted p-value are noted on top of each graph. B,F,J) Venn diagrams showing the number of overlapping statistically significant DEGs for both contrasts shown in A,E,I). Lists of overlapping genes are listed in Suppl. Table 1. C,G,K) Top 50 DEGs according to Log_2_ Fold Change for contrasts shown in A,E,I). Coloured dots mark genes involved in innate immunity-, cytokine- and interleukin-signalling according to *Panther* Gene List Analysis (Mi et al. 2017) and *GeneCards* database (Stelzer et al. 2016). D,H,L) Statistical Overrepresentation Analysis of all statistically significant DEGs for contrasts shown in A,E,I). Analysis was performed with *Panther* Reactome Pathway Analysis. Dotted line indicates Fold Enrichment Score of 1. Top three enriched pathways are depicted according to highest statistical significance.

In regulatory CD4^+^ T-cells, fewer DEGs were observed (FIG. 4E) which those which were significantly downregulated after MPXV exposure and productive infection were associated with cytokine, interleukin, and antigen-processing pathways, including *STAT1* and *IRF1*. For cycling NK cells, genes related to IFN signalling pathways were downregulated in MPXV-exposed versus mock-exposed PBMCs, but not in MPXV RNA-positive versus RNA-negative cells (FIG. 4K, L).

### Antivirals Inhibit MPXV Replication and Release of Infectious Virions in PBMCs Infected *Ex Vivo*

The efficacy of antiviral drugs against MPXV has only limitedly been studied in human cell lines (Bojkova et al. 2023). We conducted exploratory testing of two antiviral agents in unstimulated and IL-2/PHA-stimulated PBMC cultures as described above. MPXV viral protein abundance and percentage of MPXV antigen positive cells decreased after treatment with Tecovirimat and Cidofovir as shown by immunoblot (FIG. 5 A,B) and flow cytometry (Fig. 5 C,D), respectively. Both drugs effectively inhibited viral release and/or infectivity at the concentrations used, as demonstrated by plaque assays of supernatants from infected PBMCs (FIG 5 E,F).

**FIG. 5.**
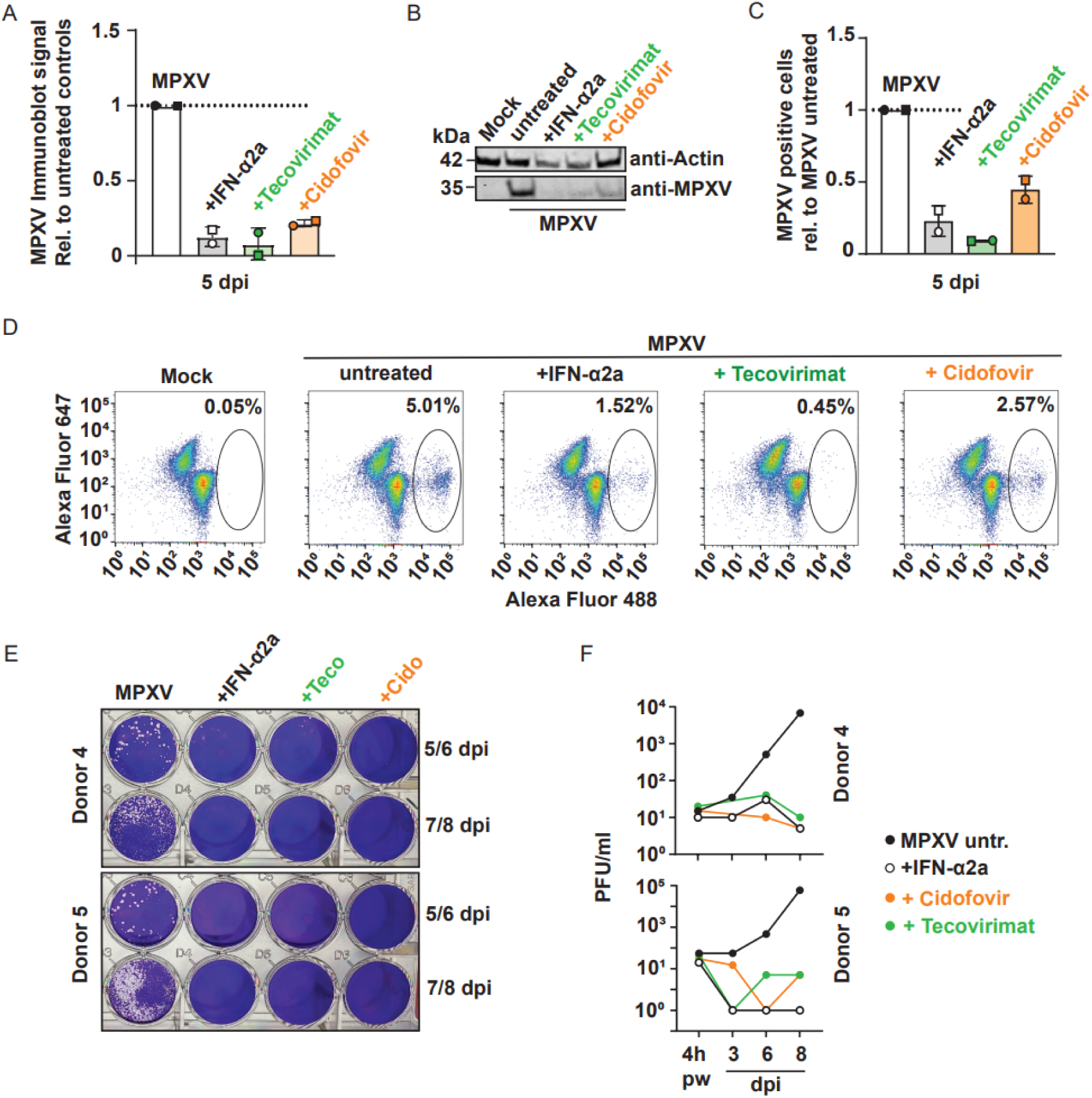
Antivirals Inhibit MPXV Replication and Release of Infectious Virions in PBMCs Infected *Ex Vivo*. PBMCs from two healthy donors were either left untreated or treated Tecovirimat (Teco, 5μM) or Cidofovir (Cido, 17μM), and exposed to MPXV (MOI 0.0006). A) Quantification of MPXV proteins in immunoblots from PBMC lysates five days post-infection. Results from two donors are presented with the mean and standard deviation. B) Representative immunoblot of MPXV proteins in lysates of PBMCs using an anti-Orthopox polyclonal rabbit serum. C) Percentage of MPXV antigen-positive cells was quantified by immunostaining of permeabilised cells with an anti-Orthopox polyclonal rabbit serum and flow cytometric analysis five days post-infection. Results from two donors are presented with the mean and standard deviation. Representative dot plots from one donor are shown in D). E) Plaque assays on Vero E6 cells from supernatants from MPXV-infected PBMCs. Supernatants were harvested at indicated time points post infection including one condition 4 hours post infection directly after removal of inoculum and washing of PBMCs with RPMI medium. F) MPXV titers from supernatants from MPXV-infected PBMC according to E).

## Discussion

Our study demonstrates susceptibility of human PBMCs to *ex vivo* infection by a MPXV clade IIb isolate and their ability to produce infectious virus progeny. Within PBMCs, monocytes, regulatory CD4^+^ T-cells and cycling NK cells appear to be the key target cells of MPXV infection. Additionally, we made first attempts to elucidate the cellular innate immune responses of individual leukocyte types to MPXV infection and demonstrate the suitability of PBMCs to evaluate antiviral drugs.

Monocytes appear to be the first primary target cells of the MPXV clade IIb, followed by delayed infection of regulatory CD4^+^ T-cells and cycling NK cells. An earlier study analysing the tropism of GFP-expressing vaccinia virus in human PBMCs identified the monocyte population as the most frequently infected cell type, followed by other immune cells (Sánchez-Puig et al. 2004). Another study (Zaucha et al. 2001) demonstrated positivity for poxviral antigens in monocytic cells from necropsies of cynomolgus monkeys that had been exposed to aerosolised MPXV clade I. Since these poxvirus antigen-positive monocytes were present in pulmonary and mediastinal lymphatics, the authors suggested that monocytes might be the primary vehicle for lymphogenous and subsequent hematogenous dissemination (Zaucha et al. 2001). This was supported by data for variola virus infection in cynomolgus monkeys, which exhibited a monocyte-associated viremia (Jahrling et al. 2004). Our results suggest a key role for monocytes as target cells for MPXV in humans, warranting more detailed research into their role in intra-host dissemination of MPXV in humans *in vivo*.

During the 2022 MPXV outbreak, individuals in the MSM community were at a higher risk of acquiring MPXV infections (Lum et al. 2022) while also carrying a disproportionate burden of HIV in European and North American countries (Lewis and Wilson 2017). Given the apparently overlapping target cell profile among PBMCs for HIV-1 and MPXV clade IIb, it is imperative to investigate a potential functional interaction of both viruses in monocytes and regulatory CD4^+^ T-cells. Monocytes represent a long-lived arm of the HIV-1 reservoir given that upon differentiation in macrophages, they can produce and archive HIV-1 for a prolonged period of time without dying (Sharova et al. 2005). Among CD4^+^ T-cells, regulatory CD4^+^ T-cells have been shown to be part of the HIV-1 reservoir under ART (Pardons et al. 2019), at equal or even higher frequencies compared to other CD4^+^ T-cell subtypes (Dunay et al. 2017; Jiao et al. 2015; Tran et al. 2008), and they may be enriched for intact, replication-competent HIV-1 proviruses as compared other CD4^+^ T-cell subsets (Pardons et al. 2019). While simultaneous acquisition of HIV-1 and MPXV might occur on rare occasions, the more relevant clinical scenario may be a potential modulation of the transcriptional status of integrated HIV-1 proviruses in people living with HIV (PLHIV) by MPXV infection. Interestingly, in the (to our knowledge) single study that monitored HIV-1 viremia pre-, during and post MPXV infection (Raccagni et al. 2023), two out of 28 PLHIV displayed increased viremia (from undetectable to 196 copies/ml, and from 263 to 1220 copies/ml) at the time point of MPOX diagnosis. Given our preliminary data showing a massive transcriptional reprogramming and pronounced downregulation of expression of genes involved in immune defence and IFN signalling by MPXV infection, we hypothesise that reversal of transcriptional HIV-1 quiescence is facilitated. Importantly, HIV-1 reactivation would not be prevented by antiretroviral treatment (ART) and may have negative impact for PLHIV, as viral gene expression is seen as a major contributor to chronic inflammation, immune activation and immunosenescence during ART (Fombellida-Lopez et al. 2024). Interestingly, in our experiments, at day three post-infection, only monocytes displayed notable quantities of viral RNA, suggesting that infection of this cell type occurs more rapidly and/or more efficiently compared to CD4^+^ T-cells and NK cells which both scored positive only at day five post-infection. An alternative explanation could be preferential cell-to-cell transmission of MPXV from infected monocytes to CD4^+^ T-cells and/or NK cells. Further experiments are required to inform whether interaction of the different cell types is essential to enable infection and decipher if all three cell types contribute equally to virus production.

MPXV downregulated the innate immune response in infected human PBMCs, particularly impacting pathways associated with IFN, interleukin, and cytokine signalling, in line with the well-documented capacity of different poxviruses to inhibit innate immunity at various levels (H. Li et al. 2023; Hernaez and Alcamí 2024; Saghazadeh and Rezaei 2022). Mechanisms include evasion of DNA-driven signalling, interference with IFN signalling, inhibition of NF-κB activation, and suppression of apoptosis (Hernaez and Alcamí 2024). For instance, in rhesus macaques infected with MPXV clade I, MPXV impaired chemokine receptor expression in NK cells and reduced IFN-γ secretion (Song et al. 2013). Additionally, MPXV encodes secreted IFN-α/β-binding proteins that block IFN interactions with cellular receptors and can evade antiviral CD8^+^ and CD4^+^ T cell responses through alternative antigen presentation (Hernáez et al. 2018; Fernández de Marco et al. 2010). It will be important to define if MPXV clades I and II and subclades differ in their ability to interfere with cell-intrinsic immunity in PBMCs, with potential implications on the efficiency of systemic dissemination.

In addition to shedding light on the cellular tropism of MPXV, we demonstrate that PBMCs are a suitable primary *ex vivo* model for evaluating antiviral compounds against MPXV in a human system. Tecovirimat, Cidofovir, and Brincidofovir are currently the only available antivirals against MPOX (Lum et al. 2022), underscoring the urgent need for intensified drug development and testing. Since these antivirals were originally developed for treatment of smallpox, they have not been extensively evaluated against MPXV. The *in vitro* efficacy of Tecovirimat and Cidofovir has been demonstrated in MPXV-infected African Green monkey VeroE6 cells (Nunes et al. 2023; Frenois-Veyrat et al. 2022; Warner et al. 2022), and only limited data exist regarding *in vitro* or *ex vivo* drug efficacy in a human model. Two studies assessed their efficacy in human primary fibroblasts, keratinocytes and skin or kidney organoids, focusing on transmission and primary infection (P. Li, Pachis, et al. 2023; Bojkova et al. 2023; P. Li, Du, et al. 2023). A PBMC-based model may not only predict the ability of drug candidates to inhibit infection but potentially also to contribute to slow down intra-host spread if effective in leukocytes, a cell type population which may not only contribute to viral production but in addition may serve as a vehicle for virus particles.

Our work has some limitations. Firstly, although our *ex vivo* model provides a standardised and reproducible method for investigating MPXV-provoked immune responses, it is not suited to capture the complexity of virus:host interactions occurring *in vivo* during disease progression. Analysing patient biomaterial instead of *ex vivo*-infected PBMCs from healthy donors will provide the full picture of merged responses mounted by the different components of the immune system during disease, as opposed to the mere cell-intrinsic response to virus infection. However, studying early dissemination *in vivo* presents a huge challenge, as it occurs before clinical symptoms become apparent, making it difficult to observe in infected individuals. In addition, in order to dissect infection events occurring at transmission sites versus systemic dissemination of the virus, and to develop therapeutic approaches which ideally interfere with both, dedicated models are indispensable. Additionally, the number of donors in our study was limited, with scRNA-seq conducted so far in PBMCs from only one donor. However, we believe that the negligible inter-individual variability in the general properties of viral tropism makes these results valuable despite this limitation.

Altogether, our study advances the understanding of MPXV infection by demonstrating that human PBMCs, particularly monocytes, regulatory CD4^+^ T cells, and cycling NK cells, are key targets for MPXV clade IIb. Our work also demonstrates the utility of PBMCs as an *ex vivo* human model for testing antiviral compounds, in times of need for further research and drug development due to limited current treatment options. Our findings provide valuable insights into a potential key mechanism of MPXV immunopathogenesis and pave the way for potential therapeutic approaches targeting systemic dissemination of MPXV.

## Acknowledgments

We thank SIGA Technologies for providing Tecovirimat.

## Author Contributions

LBJ, CG designed research.

LBJ, JJ, JM performed research.

LBJ, DP analysed data.

VC, CG supervised research, reviewed and commented on the manuscript.

LBJ, CG wrote the paper.

