## Supplement for "MPXV Infects Human PBMCs in a Type I Interferon-Sensitive Manner"

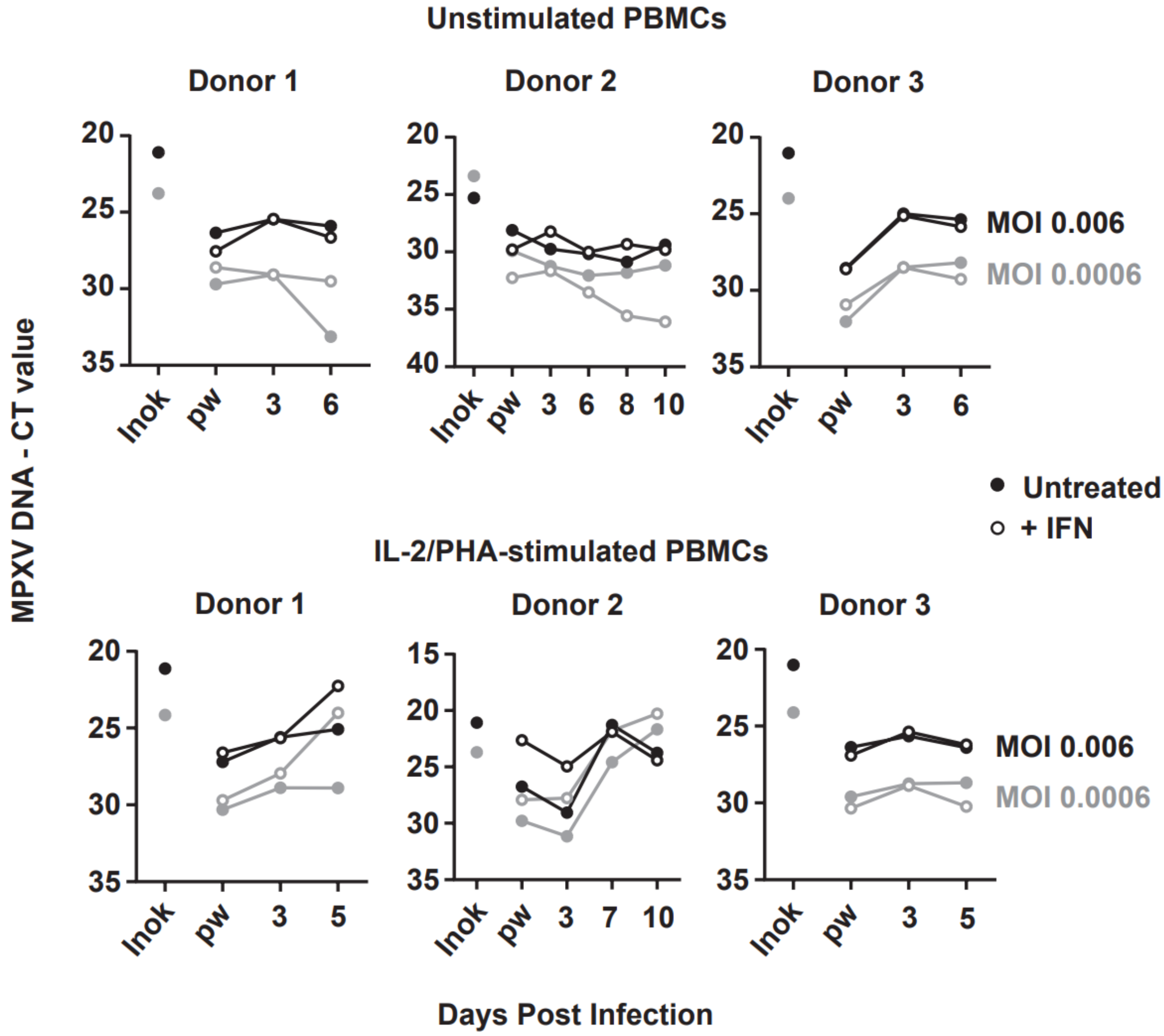

**Suppl. FIG. 1 MPXV DNA Levels in Supernatants from Infected PBMCs**

PBMCs from three healthy donors were left unstimulated (upper panels) or stimulated with IL-2/PHA (bottom panels) and exposed to MPXV at increasing MOIs in presence or absence of IFN-α2a (500 IU/ml) and supernatants were harvested at indicated time points post-infection for MPXV DNA qPCR-based quantification. Shown are the CT values of MPXV DNA.

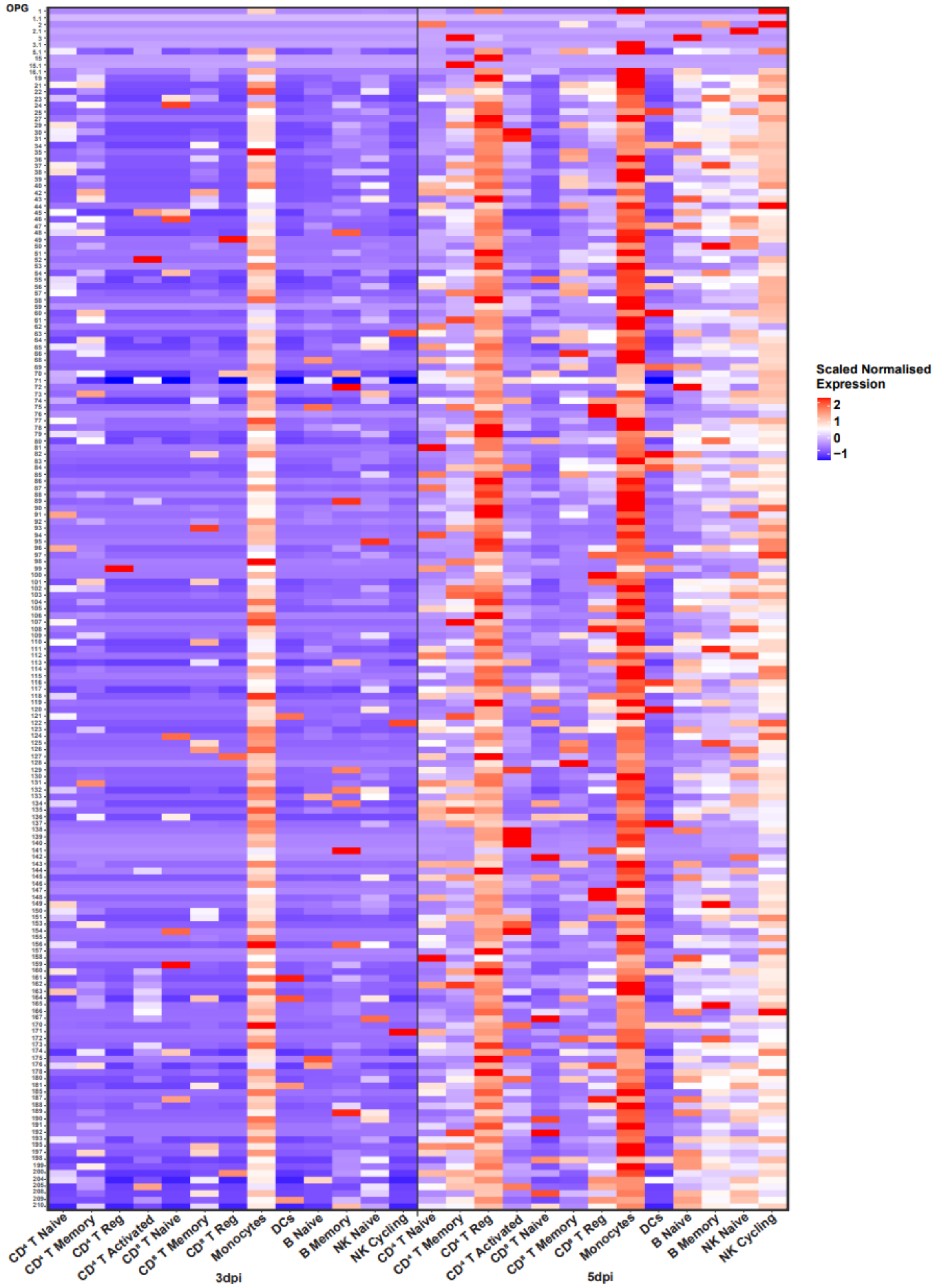

**Suppl. FIG. 2 Cell Type-Specific and Time-Dependent Expression of MPXV Genes**

The scRNA-seq dataset shown in Fig. 3 was analysed for viral gene expression in individual cell types and subsets. Shown are the scaled, normalised expression values of MPXV genes, arranged 5’ to 3’ across the viral reference genome, for MPXV-infected cells at day three (left) and five (right) post-infection.

**Suppl. Table 1 Overlapping DEGs from MPXV-Exposed vs. Mock-Exposed and MPXV RNA-Positive vs. RNA-Negative Cells**

List of overlapping statistically significant DEGs shown in Venn diagrams of FIG. 3 B,F,J. DEG analysis was performed for the contrasts MPXV-exposed vs. mock-exposed cells, and MPXV-RNA-positive cells vs. MPXV RNA-negative cells within the MPXV-exposed culture.

| **Monocytes** | **Monocytes** | **CD4 T Reg** | **CD4 T Reg** | **NK Cycling** | **NK Cycling** |
|---|---|---|---|---|---|
| **Up/down-regulated** | **Gene** | **Up/down-regulated** | **Gene** | **Up/down-regulated** | **Gene** |
| down | A2M | down | APRT | none | none |
| down | AC015660.2 | down | CFL1 | | |
| down | AC067751.1 | down | CD69 | | |
| up | AL627171.2 | down | CRIP1 | | |
| down | ALDH1A2 | down | EIF5A | | |
| down | ALCAM | down | EMP3 | | |
| down | ANXA11 | down | ENO1 | | |
| down | APRT | down | GBP2 | | |
| down | ARAP2 | down | GBP4 | | |
| down | ARL6IP5 | down | IL32 | | |

| Monocytes | Monocytes | | |
|---|---|---|---|
| **Up/down-regulated** | **Gene** | | |
| down | ATP5F1B | down | IRF1 |
| down | ATP5F1D | down | LY6E |
| down | ATP6V0E1 | down | MIR155HG |
| down | B2M | down | MYL12A |
| down | BCL2A1 | down | MYL12B |
| down | BTN3A2 | down | PARP14 |
| down | CALM3 | down | PKM |
| down | CD274 | down | PSMB3 |
| down | CD52 | down | PSMB8 |
| down | CD53 | down | PSMB9 |
| down | CD74 | down | PSME1 |
| down | CD83 | down | PSME2 |
| down | CD86 | down | RNASEK |
| down | COX5B | down | S100A10 |
| down | COX6A1 | down | S100A11 |

| **Monocytes** | **Monocytes** | | |
|---|---|---|---|
| **Up/down-regulated** | **Gene** | | |
| down | COX8A | up | SRPK2 |
| down | CRABP2 | down | TAP1 |
| down | CRIP1 | down | TXN |
| down | C19orf53 | | |
| down | C15orf48 | | |
| down | CSF1 | | |
| down | CYP27B1 | | |
| down | EPSTI1 | | |
| down | FABP5 | | |
| down | FERMT3 | | |
| down | FKBP1A | | |
| down | FGL2 | | |
| down | GBP1 | | |
| down | GBP2 | | |
| down | GBP3 | | |
| down | GBP4 | | |

| **Monocytes** | **Monocytes** |
|---|---|
| **Up/down-regulated** | **Gene** |
| down | GBP5 |
| down | GSTO1 |
| down | GSN |
| down | HLA-A |
| down | HLA-B |
| down | HLA-C |
| down | HLA-DMA |
| down | HLA-DPA1 |
| down | HLA-DPB1 |
| down | HLA-DQA1 |
| down | HLA-DRB1 |
| down | HLA-DRB5 |
| down | HLA-E |
| down | HLA-F |
| down | ICAM1 |
| down | IDO1 |
| down | IFI35 |

| Monocytes | Monocytes |
|---|---|
| **Up/down-regulated** | **Gene** |
| down | IFI6 |
| down | IFIT3 |
| down | IGFBP6 |
| down | IL1RN |
| down | IL2RG |
| down | JAK2 |
| down | LY6E |
| down | MGLL |
| down | MVP |
| down | MYL6 |
| down | MYL12A |
| down | NLRC5 |
| down | N4BP2L1 |
| down | NIBAN1 |
| down | NINJ1 |
| down | NPC2 |

| Monocytes | Monocytes |
|---|---|
| **Up/down-regulated** | **Gene** |
| down | NRP2 |
| down | OAS1 |
| down | PARP9 |
| down | PARP14 |
| down | PET100 |
| down | PKM |
| down | PRDX5 |
| down | PSENEN |
| down | PSMB9 |
| down | PSTPIP2 |
| down | RNASEK |
| down | SCIMP |
| down | SLAMF7 |
| down | SLC1A2 |
| down | SLC15A3 |
| down | SLC31A2 |

| **Monocytes** | **Monocytes** |
|---|---|
| **Up/down-regulated** | **Gene** |
| down | SQOR |
| down | SRGN |
| down | SSR2 |
| down | STAT1 |
| down | TGM2 |
| down | TIMM8B |
| down | TMEM14C |
| down | TMSB10 |
| down | TNIP1 |
| down | TRAF1 |
| down | TRIM69 |
| down | TRIR |
| down | TXN |
| down | UBE2L6 |
| down | ZBTB38 |
